# Tadpoles develop elevated heat tolerance in urban heat islands regardless of sex

**DOI:** 10.1101/2023.08.26.554938

**Authors:** Veronika Bókony, Emese Balogh, János Ujszegi, Nikolett Ujhegyi, Márk Szederkényi, Attila Hettyey

## Abstract

The ability of wildlife to endure the effects of high temperatures is increasingly important for biodiversity conservation under climate change and spreading urbanization. Organisms living in urban heat islands can have elevated heat tolerance *via* both phenotypic plasticity and microevolution. However, the prevalence and mechanisms of such thermal adaptations are barely known in aquatic organisms. Furthermore, males and females can differ in heat tolerance, which may lead to sex-biased mortality, yet it is unknown how sex differences in thermal biology influence urban adaptations. To address these knowledge gaps, we measured critical thermal maxima (CT_max_) in male and female agile frog (*Rana dalmatina*) tadpoles captured from warm urban ponds and cool woodland ponds, and in a common-garden experiment where embryos collected from both habitat types were raised in the laboratory. We found higher CT_max_ in urban-dwelling tadpoles compared to their counterparts living in woodland ponds. This difference was reversed in the common-garden experiment: tadpoles originating from urban ponds had lower CT_max_ than tadpoles originating from woodland ponds. We found no effect of sex on CT_max_ or its difference between habitats. These results demonstrate that aquatic amphibian larvae can respond to the urban heat island effect with increased heat tolerance similarly to the other, mostly terrestrial taxa studied so far, and that phenotypic plasticity may be the main driver of this response. Our findings also suggest that heat-induced mortality may be independent of sex in tadpoles, but research is needed in many more taxa to explore potentially sex-dependent urban thermal responses.

## Introduction

Heat tolerance, i.e. the capacity to cope with high temperatures, is becoming increasingly important throughout the tree of life with ongoing climate change. In many regions across the globe, not only are average temperatures rising but heat events are also getting more frequent (Perkins-Kirkpatrick & Lewis, 2020). High temperature can be directly lethal by physiological breakdown due to heat stress, but it may also cause secondary mortality by making organisms more susceptible to predation or disease (Kroeker & Sanford, 2022). Populations can adapt to better tolerate high temperatures by two mutually non-exclusive mechanisms: phenotypic plasticity expressed during the lifetime of individuals (acclimation or thermal plasticity), or microevolutionary and epigenetic changes over generations (Urban *et al*., 2014). Mechanisms matter because phenotypically plastic responses can manifest much faster but can also be costly and their scope may be constrained (Gunderson & Stillman, 2015; Murren *et al*., 2015; Radchuk *et al*., 2019).

Dealing with heat stress is especially relevant for organisms living in urbanized habitats, because heat storage in buildings and sealed roads makes cities warmer compared to surrounding non-urban areas, and this urban heat island effect is amplified during heat waves (Li & Bou-Zeid, 2013). Accordingly, it has been shown in a variety of ectothermic species that urban populations have higher heat tolerance, expressed as the critical thermal maximum (CT_max_), the upper temperature at which animals lose the ability to function (Diamond & Martin, 2021). These phenotypic changes can be adaptive (Brans *et al*., 2017; Martin *et al*., 2021) and result from a combination of phenotypic plasticity and microevolution (Brans *et al*., 2017; Diamond *et al*., 2017).

Heat tolerance may also differ between sexes. In humans, for example, females are at higher risk of dying than males during heat waves (van Steen *et al*., 2019). This has important implications for evolutionary ecology and conservation biology. Sex-dependent mortality can lead to skewed sex ratios, which then constrain effective population size and adaptive potential but can also catalyze evolutionary changes in sex-specific life histories and social systems (Mitchell & Janzen, 2010; Schacht *et al*., 2022). Also, skewed sex ratios can have cascading effects on other species and even ecosystems (Edmands, 2021). Despite this significance of the issue, there is a dearth of information on sex differences in heat tolerance (Edmands, 2021; Pottier *et al*., 2021). Furthermore, sexes may also differ in their capacity for thermal plasticity (Pottier *et al*., 2021), yet next to nothing is known about the role of sex in thermal adaptations to urban heat islands.

For several organismal traits such as hormone levels, cognitive performance, and parasite load, it has been demonstrated that urbanization can have sex-specific effects (Bonier *et al*., 2007; Preiszner *et al*., 2017; Sykes *et al*., 2021). Because males and females can differ in the temperatures they experience, prefer, or tolerate due to differences in size, life history, and behaviors associated with reproduction (Ruckstuhl & Neuhaus, 2005; Edmands, 2021), the sexes may also differ in the selection pressures that urban heat islands exert on them. Then, sex-dependent selection for thermal tolerance and/or for acclimation capacity may complicate thermal adaptation of urban populations due to genetic correlations between the sexes and sexual selection (Leith *et al*., 2022). However, our understanding of these potential outcomes is highly deficient due to the virtual lack of empirical studies on sex differences in the effects of urbanization on heat tolerance.

Although the number of studies published on urban heat tolerance is increasing exponentially (Roeder *et al*., 2021), almost all this effort has been focused on terrestrial organisms (Diamond & Martin, 2021). Aquatic ectotherms, however, are also exposed to the urban heat island effect (Brans *et al*., 2018) and may be especially vulnerable to it due to limited dispersal (Pagliaro & Knouft, 2020) and the conflict between increased demand for and decreased supply of oxygen in warmer water (Brans *et al*., 2017). Here, we investigated heat tolerance in tadpoles of the agile frog (*Rana dalmatina*), a European species with decreasing population trends that occurs in both urban and non-urban habitats (Kaya *et al*., 2009). First, we show that individuals from relatively warm urban ponds and those from relatively cool non-urban ponds differ in heat tolerance measured as CT_max_. Second, we report a common garden experiment to infer whether this difference is attributable to individual plasticity or transgenerational change. Finally, we compare heat tolerance between males and females and test whether the effect of urbanization on heat tolerance is sex-dependent.

## Methods

### Pond temperatures

We used six study sites: three were in hilly woodlands with <0.1% anthropogenically modified habitat within 500 m of each pond, whereas the other three sites were in three different townships with ca. 70% anthropogenically modified land cover within 500 m of the ponds (Table S1). In each pond, we recorded water temperature every 30 minutes from 6^th^ April to 8^th^ July 2022 using Onset UA-002-64 HOBO loggers. We placed four loggers within each pond in the area where we collected eggs (see below). One pair of loggers was placed in the deepest water we could access, whereas another pair of loggers was placed close to the shore, at ≤ 30 cm water depth. Within each pair, one logger was ca. 5 cm under the water surface and one was ca. 5 cm above the bottom. We monitored the loggers weekly and adjusted their position to follow changes in water depth (for further information see Fig. S1).

### Experimental protocol

This study was approved by the Ethics Committee of the Plant Protection Institute and licensed by the Environment Protection and Nature Conservation Department of the Pest County Bureau of the Hungarian Government (PE-06/KTF/00754-8/2022, PE-06/KTF/00754- 9/2022, PE-06/KTF/00754-10/2022, PE/EA/295-7/2018). From the six study ponds, we collected three cohorts of animals. For the first two cohorts, we collected freshly spawned agile frog eggs on 4^th^ April, and three-weeks old embryos (right before hatching) on 20^th^ April. From each pond, we took ca. 20 embryos from each of four egg masses for each of the two cohorts (Table S1) and transported them to our laboratory. We kept each sibling group in a separate container with ca. 1 cm deep reconstituted soft water (RSW; 48 mg NaHCO_3_, 30 mg CaSO_4_ × 2 H_2_O, 61 mg MgSO_4_ × 7 H_2_O, 2 mg KCl added to 1 L reverse-osmosis filtered, UV-sterilized, aerated tap water). Over the course of the study, temperature in the lab was set to gradually increase from 18 °C to 20 °C (mean ± standard deviation: 19.3 ± 0.9 °C) and we regularly adjusted the photoperiod to mimic the natural dark-light cycles. When the animals reached the free-swimming state, i.e. developmental stage 25 according to (Gosner, 1960), we placed them individually in 2-L plastic rearing containers filled with 1 L RSW, arranged in a randomized block design to ensure that both cohorts and all six populations were homogeneously distributed across the shelves in the laboratory. We changed the rearing water twice a week and fed the tadpoles *ad libitum* with chopped, slightly boiled spinach. On the 18^th^ - 20^th^ day after reaching developmental stage 25, we randomly selected six tadpoles from each sibling group (Table S1), resulting in a total sample size of 144 tadpoles (24 per pond) from each cohort, and tested their CT_max_ (see below). The remaining tadpoles were released at their ponds of origin.

From the same six ponds, we collected the third cohort as tadpoles by dip-netting in the second half of May, when the animals were at a similar developmental stage (having only small hindlimb buds) as the captive-reared tadpoles were at the time of CT_max_ testing. We aimed to collect 24 tadpoles from each pond, but we did not find any in one of the urban ponds and we could capture only six from a woodland (non-urban) pond, yielding a total sample size of 102 free-living tadpoles (Table S1). We transported the tadpoles to our laboratory, housed and fed them the same way as the captive-reared tadpoles, and tested their CT_max_ one or two days after their collection.

We measured CT_max_ in a randomized order within each cohort by placing 8 tadpoles, each in its original rearing container, into a tray (80 × 60 × 12 cm) in which water was heated with two digital thermostat heaters (BRH Heizung LCD Turbo 600) and circulated by two water pumps (Tetra WP 300). For the time of the test, each tadpole’s container was filled with 1.5 L fresh RSW, and when all 8 containers were placed in the tray, water level in the tray was ca. 0.5 cm below that in the tadpole containers. We increased water temperature in the tadpoles’ containers at a rate of 0.6°C/min. Twenty-one minutes after placing the containers into the tray, we started to observe the animals, lightly prodding the base of their tail every 6 seconds. We defined CT_max_ as the temperature (as measured by Greisinger digital thermometers GTH175/PT; ± 0.1 °C) at which the tadpole failed to respond with motion over three consecutive prods. The test was performed by six experimenters, four at a time, each person overseeing two tadpoles always at the same two positions within the tray.

After the CT_max_ test, we weighed each tadpole (± 0.1 mg), recorded its developmental stage by stereomicroscopic examination, and stored the euthanized animals in 96% ethanol. We extracted DNA using E.Z.N.A. Tissue DNA Kit following the manufacturer’s protocol, except that digestion time was at least 3 hours. For genetic sexing, we used the method of Nemesházi *et al*. (2020). Briefly, we tested all tadpoles for sex marker Rds3 (≥ 95% sex linkage; primers: Rds3-HRM-F and Rds3-HRM-R) using high-resolution melting (Fig. S2). The total HRM reaction volume was 15 μl, containing 7.5 μl 2x PerfeCTa® SYBR® Green SuperMixes (ROX, Quantabio), 1 μl forward and 1 μl reverse primer (10 μM each), and 80- 100 ng genomic DNA in MQ water to reach the final volume. Reactions were performed in a Quantabio Q 4-channel qPCR Instrument and the results were analysed with the 1.0.2. version Q-qPCR Software (Quantabio).

### Statistical analyses

We used R 4.2.2 for all analyses (R Core Team, 2022). We analyzed pond temperatures using a generalized additive mixed model (‘gamm’ function of package ‘mgcv), because the change of temperature over time was not linear (Fig. S1). We included pond identity and logger position as fixed factors, and time as a covariate and temporal autocorrelation (order-1 auto-regressive model) within the data of each logger. For comparison among the ponds, we extracted marginal means for each pond from the model and calculated linear contrasts pairwise and also between the three urban and three woodland ponds (‘emmeans’ function of package ‘emmeans’). For the pairwise comparisons, we corrected the P-values with the false discovery rate (FDR) method (Pike, 2011).

To analyze CT_max_, we used a generalized estimation equations (GEE) model (‘geeglm’ function of package ‘geepack’). GEE is a population-averaging method that can handle the correlation structure of our data (i.e. tadpoles from the same pond are not independent, but the pond effect is nested within the habitat effect) appropriately and without penalizing power (Zuur *et al*., 2009). We included the following fixed factors: habitat type (urban or woodland), cohort (collected as eggs, embryos, or tadpoles), sex, their three-way and all two-way interactions, body mass, developmental stage, and experimenter identity; we used pond of origin as a random factor (‘compound symmetry’ correlation structure). From this initial full model, we evaluated the significance of interactions by type-2 analysis-of-deviance tables. Then we dropped the non-significant interactions stepwise, to facilitate the accuracy and ease of estimation for the significant interaction term. We calculated linear contrasts (as above, correcting the P-values with the FDR method) to test the habitat effect within each cohort from the final GEE model that contained only significant interactions but all main effects regardless of their significance.

## Results

All urban ponds were significantly warmer than all woodland ponds (Fig. S1, Fig. S3), on average by 5.58°C ± 0.17 SE (urban-woodland contrast, P < 0.001). For CT_max_, the three-way interaction between sex, cohort, and habitat of origin was non-significant (χ^2^_2_ = 2.70, P = 0.259; Fig. S4), and so were the two-way interactions between sex and cohort (χ ^2^_2_ = 1.38, P = 0.501) and between sex and habitat of origin (χ ^2^_1_ = 0.13, P = 0.713). According to the final model, females had slightly higher CT_max_ than males (by 0.07 ± 0.04 °C), but this difference was not statistically significant, although not far from the 5% significance threshold (χ ^2^_1_ = 2.87, P = 0.090). The interaction between cohort and habitat of origin was highly significant (χ ^2^_2_ = 58.4, P < 0.001; Fig. 1), such that tadpoles living in urban habitats had higher CT_max_ than tadpoles living in woodland habitats (by 0.15 ± 0.05 °C, P = 0.004), but this difference was reversed when the animals were raised in a common, captive environment after being collected from the field as freshly spawned eggs (by 0.17 ± 0.09 °C, P = 0.0451) or three-weeks old embryos (by 0.20 ± 0.08 °C, P = 0.016).

**Figure 1.**
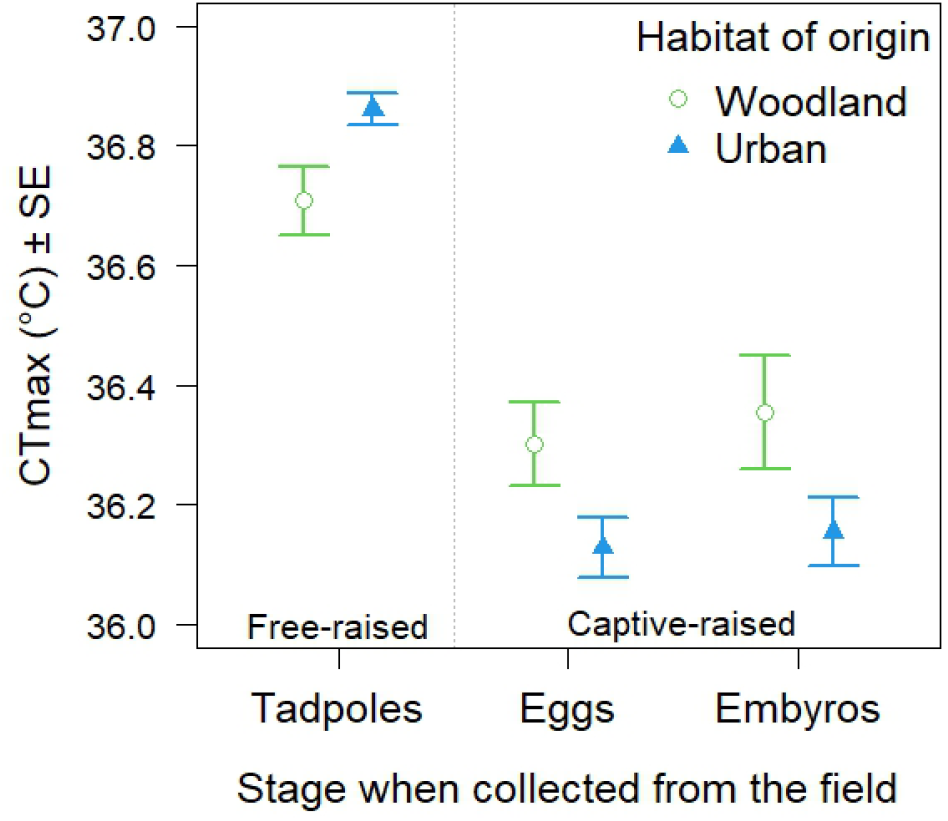
Critical thermal maximum (CT_max_) of agile frog tadpoles originating from woodland or urban habitats and raised in the field or in captivity from the egg or embryo stage. Means with standard errors (SE) were corrected for body mass, developmental stage, experimenter identity, and pseudoreplication (non-independence of tadpoles from the same site) using a generalized estimation equations model.

## Discussion

We found that urban ponds were several degrees warmer, supporting Brans *et al*. (2018) and Pagliaro & Knouft (2020) that urbanization is accompanied by higher temperatures of freshwater habitats in the temperate climate zone. This urban heat island effect was accompanied by a small but statistically significant increase in CT_max_ for agile frog tadpoles developing in urban ponds compared to their counterparts inhabiting woodland ponds. This aligns with the findings of Brans *et al*. (2017) on water fleas, demonstrating that aquatic organisms respond to urbanization by higher heat tolerance similarly to a wide variety of terrestrial taxa ranging from fungi through arthropods to lizards (McLean *et al*., 2005; Campbell-Staton *et al*., 2020; Diamond & Martin, 2021). This phenotypic response is important for several reasons. First, already these days, pond temperatures occasionally rise so high (Lambert *et al*., 2018; Lindauer *et al*., 2020) that they approach or exceed the upper thermal tolerance limits of aquatic animals (Brans *et al*., 2017; Pagliaro & Knouft, 2020), especially so in urban heat islands (see Fig. S1); and such heat events are expected to get more frequent in the future due to climate change. Second, CT_max_ is not a standalone physiological trait, as it may be part of a “thermal syndrome”, linked with other aspects of thermal performance and preference, and potentially also with behavioral and life-history traits (Goulet *et al*., 2017a; b). For example, individuals with higher CT_max_ can also better tolerate ecologically relevant slow-rate warming (Åsheim *et al*., 2020). Often, the magnitude of heat tolerance decreases with the duration of heat (Troia, 2023); for example, agile frog tadpoles can experience high mortality during a 6-days long period at 28°C (Ujszegi *et al*., 2022) despite their ≥ 33.6°C CT_max_ (according to the present study). Such a several-days heat wave has not only lethal effects but also induces sex reversal in developing larvae (Ujszegi *et al*., 2022) which may have wide-ranging consequences for individual fitness and population persistence (Bókony *et al*., 2017, 2021b; Nemesházi *et al*., 2021). Thus, a “thermal type” with elevated tolerance against the various physiological harms of high temperatures is likely to be adaptive particularly for urban populations which are more frequently exposed to heat.

This surmised adaptation can come about through individual plasticity or by changes accumulating over the course of many generations such as microevolution. Our study suggests that the former mechanism is the main driver of higher CT_max_ in urban agile frog tadpoles, because we observed elevated heat tolerance only in the free-living urban animals and not in the urban-collected animals raised in the common-garden experiment from early or late embryonic stages. Phenotypic plasticity is further suggested by the higher CT_max_ of free-living tadpoles which experienced higher temperatures in the field (up to 28.46 °C before being collected for CT_max_ testing; Fig. S1) than did their captive-raised counterparts (up to 19.9 °C). Our study was not designed to identify the mechanism of phenotypic plasticity or to measure the contribution of short-term, reversible acclimatization and longer-lasting, more slowly establishing developmental plasticity (Beaman *et al*., 2016). However, CT_max_ of amphibians can acclimate within a few days (Hutchison & Maness, 1979; Layne & Claussen, 1982; Turriago *et al*., 2023), suggesting that the 1-2 days acclimation period in our study might have canceled (some of the) differences that had been induced by acclimatization to the temperatures experienced in urban and woodland ponds, leaving room for developmental plasticity to potentially manifest in our measurements.

The lack of support for microevolution in tadpole CT_max_ is in contrast with findings on arthropods where, besides phenotypic plasticity, microevolution also proved an important driver of urban heat tolerance (Diamond & Martin, 2021). It is possible that the relatively long generation time in frogs may not have permitted enough evolutionary change to accumulate in urban populations, despite the relatively restricted gene flow across the barriers represented by anthropogenically altered habitats (Hitchings & Beebee, 1997; Lesbarrères *et al*., 2006). Also, phenotypic plasticity seems to be favored in amphibian physiological traits, maybe due to high spatiotemporal environmental heterogeneity within urban habitats (Bókony *et al*., 2019, 2021a), which is exemplified by the relatively large microclimatic differences between nearby ponds in the present study (e.g. two ponds in Pilisszentiván, see Fig. S1 & S3). Indeed, the magnitude of thermal pollution in urban ponds is strongly affected by local-scale variation such as the “park cooling effect” and runoff (Brans *et al*., 2018), which might explain why higher heat tolerance was not found in some ectotherms in broad-scale urban regions (Diamond & Martin, 2021). Naturally, heterogeneous thermal microhabitats are also present outside cities, which may select for genotypes with different thermal tolerances and thereby facilitate adaptations to urban habitats (Campbell-Staton *et al*., 2020). However, such standing variation in heat tolerance might be limited in the agile frog because it primarily occurs in relatively cool habitats and its genetic diversity across Europe is very low (Vences *et al*., 2013).

Interestingly, in our common-garden experiment, the lack of experiencing field temperatures was not simply accompanied by no difference in CT_max_; instead, the captive-raised tadpoles originating from urban ponds had lower heat tolerance than those from woodland ponds. We can only speculate about the causes of this difference. It has been demonstrated in some anuran species, including agile frogs, that compared to conspecifics originating from more natural habitats, individuals originating from urban habitats perform poorly in various fitness-related traits when raised in captivity, including slower development, reduced growth, more frequent developmental abnormalities, and higher mortality (Hitchings & Beebee, 1997; Bókony *et al*., 2018, 2023). Lower heat tolerance might be another manifestation of this overall poor physiological fitness. The reasons are unclear but may be related to transgenerational effects of endocrine-disrupting chemical pollutants (Bókony *et al*., 2018) or inbreeding in isolated urban populations due to landscape fragmentation (Hitchings & Beebee, 1997; Lesbarrères *et al*., 2006).

Finally, we found no difference between male and female tadpoles in the effect of urbanization on CT_max_, although there was a slight, non-significant trend for higher heat tolerance in females. In adult agile frogs, the two sexes may face different selection pressures on thermal tolerance for several reasons. For example, females typically mature at larger body sizes than males, which may affect multiple aspects of their thermal physiology (Rohr *et al*., 2018; Rubalcaba *et al*., 2019). Furthermore, males migrate earlier from hibernacula to the breeding ponds, sometimes even when snow and ice have not yet melted (Riis, 1997), whereas females were reported to forage less in open microhabitats compared to males (Cicort-Lucaciu *et al*., 2011). However, even if these microclimatic differences select for different thermal physiology in the two sexes, it is possible that those differences may not become expressed until sexual maturity, as sex seems to have little effect on behavior and life history in immature agile frogs (Bókony *et al*., 2021b). More studies are needed, ideally with multiple age groups and life-history stages, to increase our knowledge from next to nothing about sex differences in temperature sensitivities of amphibians and various other taxa (Edmands, 2021). That knowledge will be necessary for inferring the role of sex-biased mortality *via* sex-dependent thermal tolerance in the effects of urbanization and climate change on the fate of wildlife populations and biodiversity.

## Acknowledgements

We thank Csenge Kalina, Zsanett Mikó, and all other members of the Evolutionary Ecology Research Group for their help during the experiment. We are grateful to Bernadett Zsinka (Molecular Ecology Research Group of the University of Veterinary Medicine, Budapest) for help in the DNA work, Julia Halász (Institute of Genetics and Biotechnology, Hungarian University of Agriculture and Life Sciences, Budapest) for help in measuring DNA concentration, and Daniel Teutsch (Quantabio) for the qPCR machine calibration. The study was supported by the National Research, Development and Innovation Office of Hungary (NKFIH K-135016 & K-124375), the New National Excellence Program of the Ministry for Innovation and Technology (ÚNKP-21-5 to VB and AH, ÚNKP-22-5 to AH, ÚNKP-22-3-1 to EB, ÚNKP-22-4-II to UJ), the University of Veterinary Medicine Budapest (EB and VB), and the János Bolyai Scholarship of the Hungarian Academy of Sciences to VB and AH.

## Author contributions

- VB: planning, funding, tadpole raising, CTmax, analysis, first draft, review & editing
- EB: tadpole raising, genetic sexing, review & editing
- JU: planning, tadpole raising, CTmax, review & editing
- NU: planning, tadpole raising, CTmax, review & editing
- MS: tadpole raising, CTmax, logger management, review & editing
- AH: planning, funding, CTmax, review & editing

## Supplementary Material

**Table S1.** Study sites for water temperature measurements, and sample sizes for CTmax.

| Pond | Habitat type | Latitude | Longitude | Number of tadpoles tested for CTmax, collected as: |  |  |
| --- | --- | --- | --- | --- | --- | --- |
|  |  |  |  | Eggs<sup>*</sup> | Embryos<sup>*</sup> | Tadpoles |
| Felsőhosszúréti | Woodland | 47°43'37.00" | 19°01'02.00" | 24 | 24 | 24 |
| Mélymocsár | Woodland | 47°42'27.00" | 19°02'24.00" | 24 | 24 | 6 |
| Papréti | Woodland | 47°44'20.00" | 19°00'43.00" | 24 | 24 | 24 |
| Pilisvörösvár township<sup>†</sup> |  |  |  |  |  |  |
| Cigány-tó | Urban | 47°36'24.08" | 18°55'21.40" | 30 | 30 | 24 |
| Pálya-tó | Urban | 47°36'39.14" | 18°55'21.49" | 0 | 0 | 0 |
| Pilisszentiván township<sup>†</sup> |  |  |  |  |  |  |
| Small pond | Urban | 47°36'25.55" | 18°54'38.15" | 24 | 24 | 24 |
| Big pond | Urban | 47°36'21.89" | 18°54'32.68" |  |  |  |
| Pilisszántó township |  |  |  |  |  |  |
| Határréti-tó | Urban | 47°38'48.20" | 18°54'31.25" | 18 | 18 | 0 |

<sup>†</sup>The two ponds within each of the two townships are next to each other and therefore were treated as one site (i.e. one population of agile frogs). All 6 sites are >1 km from each other; the closest two sites (Pilisvörösvár and Pilisszentiván) are separated by roads, railroads, and buildings.
<sup>\*</sup>We sampled 4 egg masses at each pond, except that we found only 3 egg masses of appropriate age at Határréti-tó, so we added a 5th egg mass from Cigány-tó. From each egg mass within each age group, we tested 6 tadpoles for CTmax, except for two egg masses where only 4 or 5 tadpoles were in the appropriate developmental stage for testing and therefore were substituted by three tadpoles from other egg masses from the same pond.

**Figure S1.**
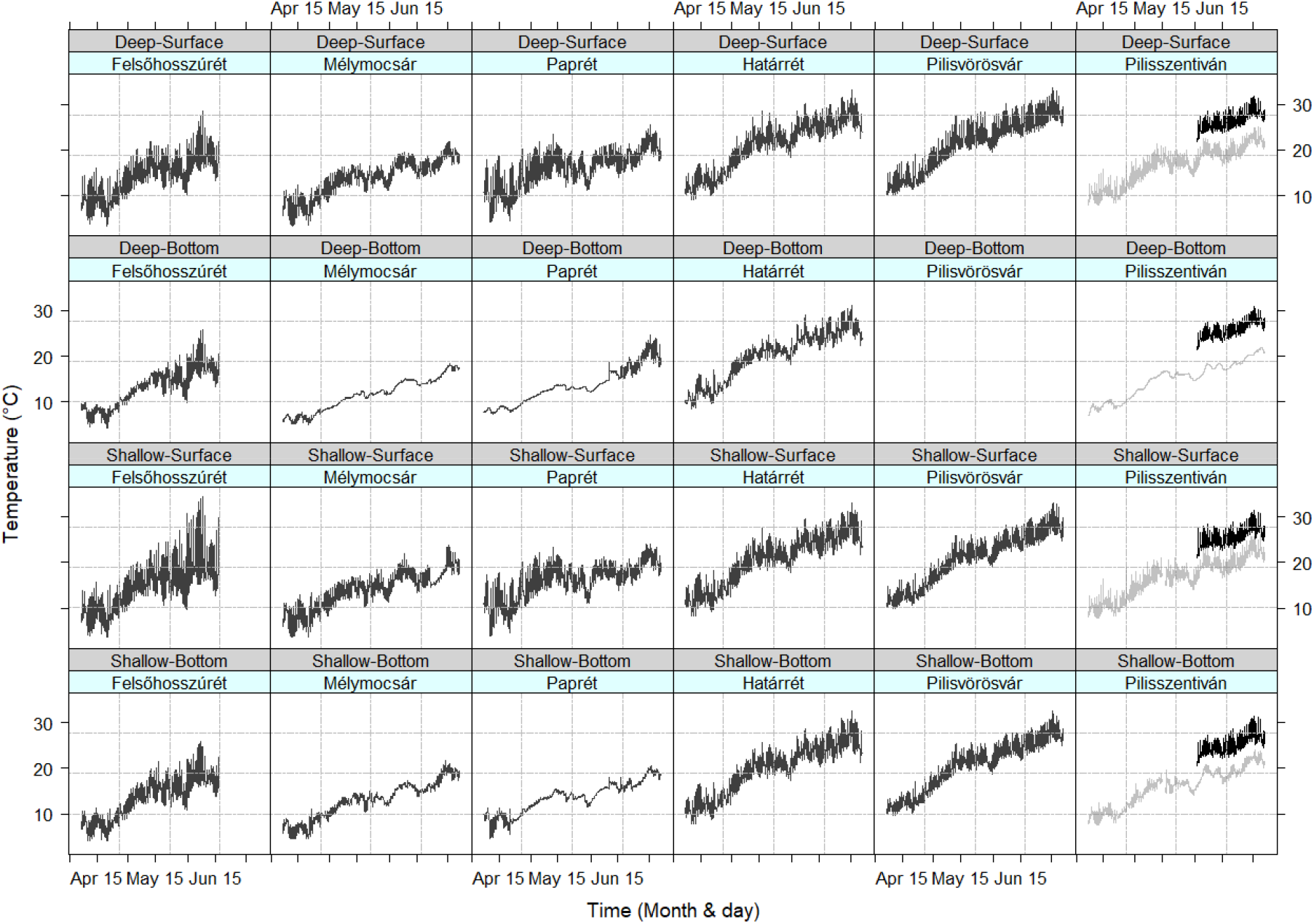
Water temperatures in 2022, measured at four locations within each pond over the tadpole development season (6 April – 8 July). The three sites on the left (Felsőhosszúrét, Mélymocsár, Paprét) are non-urban (woodland); the three sites on the right (Határrét, Pilisvörösvár, Pilisszentiván) are urban. Note that one of the non-urban ponds, Felsőhosszúrét, dried out so we have no data for this pond after 16 June. In Pilisszentiván, there are two ponds right next to each other; we measured temperatures and collected animals in both: a small, shady pond (grey lines; the gap at 15-16 May is due to drying) and a large, open pond (black lines; we have data only from 2 June due to logistic reasons).

**Figure S2.**
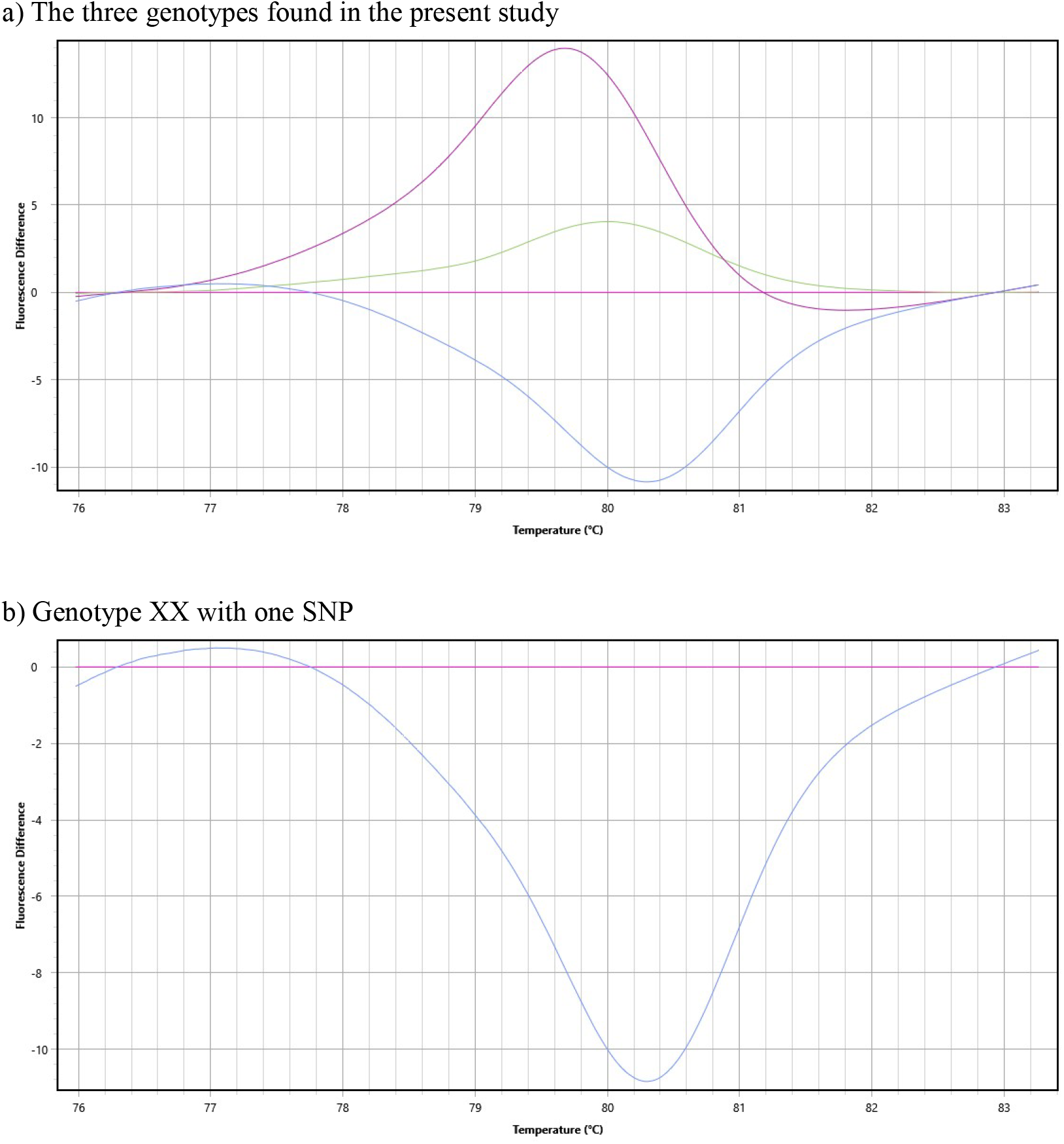
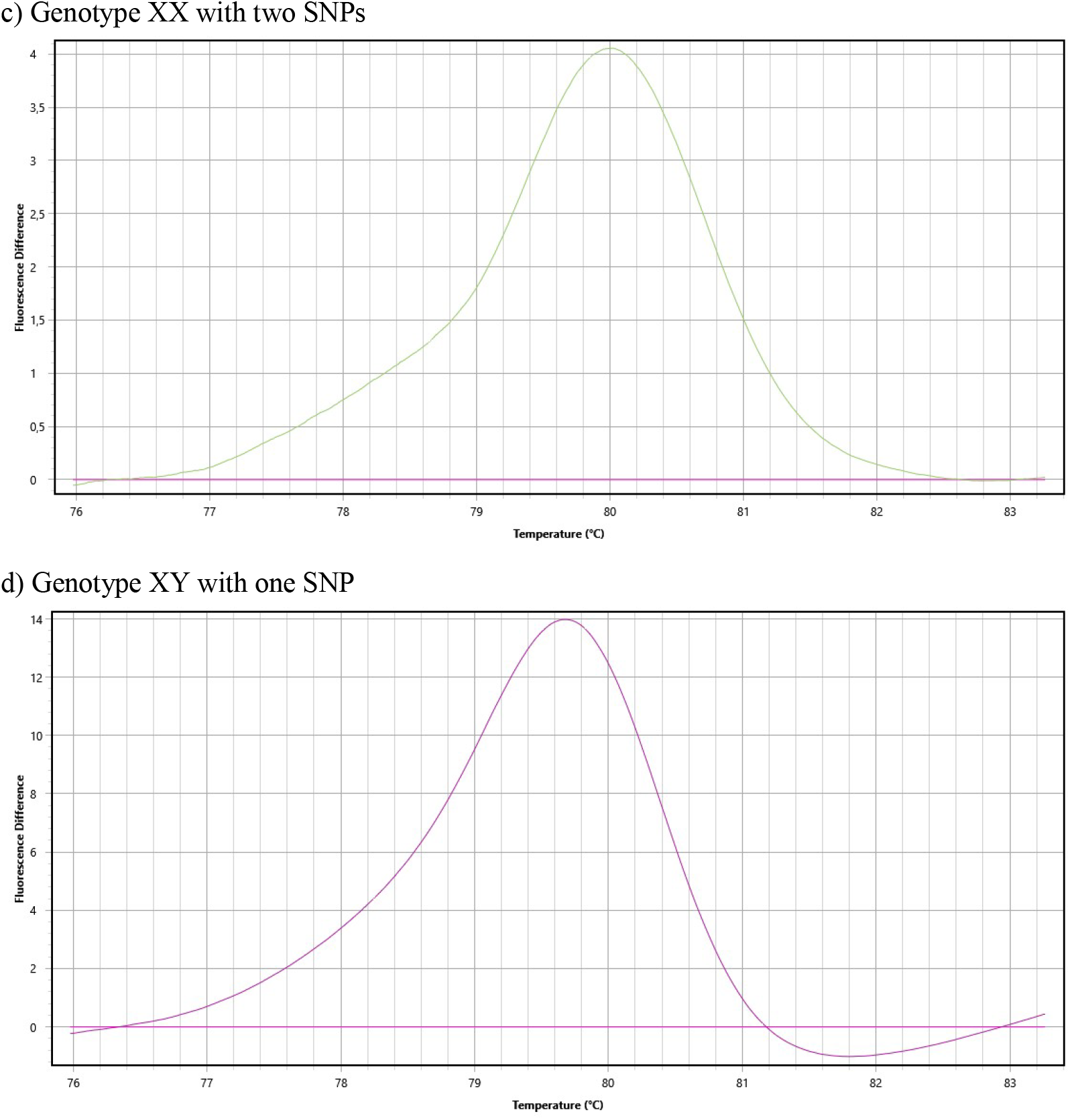
HRM-based genotyping with marker Rds3. Curves on the Difference Plots were drawn by Q-qPCR Software v. 1.0.2 (Quantabio). Besides the single nucleotide polymorphism (SNP) used for sexing, in some individuals a second SNP occurs, altering in the curves’ shape (for more details, see Fig. S2 in Nemesházi et al. 2020). In the present study, some of the genetic females (XX) but none of the genetic males (XY) had the second SNP.

**Figure S3.**
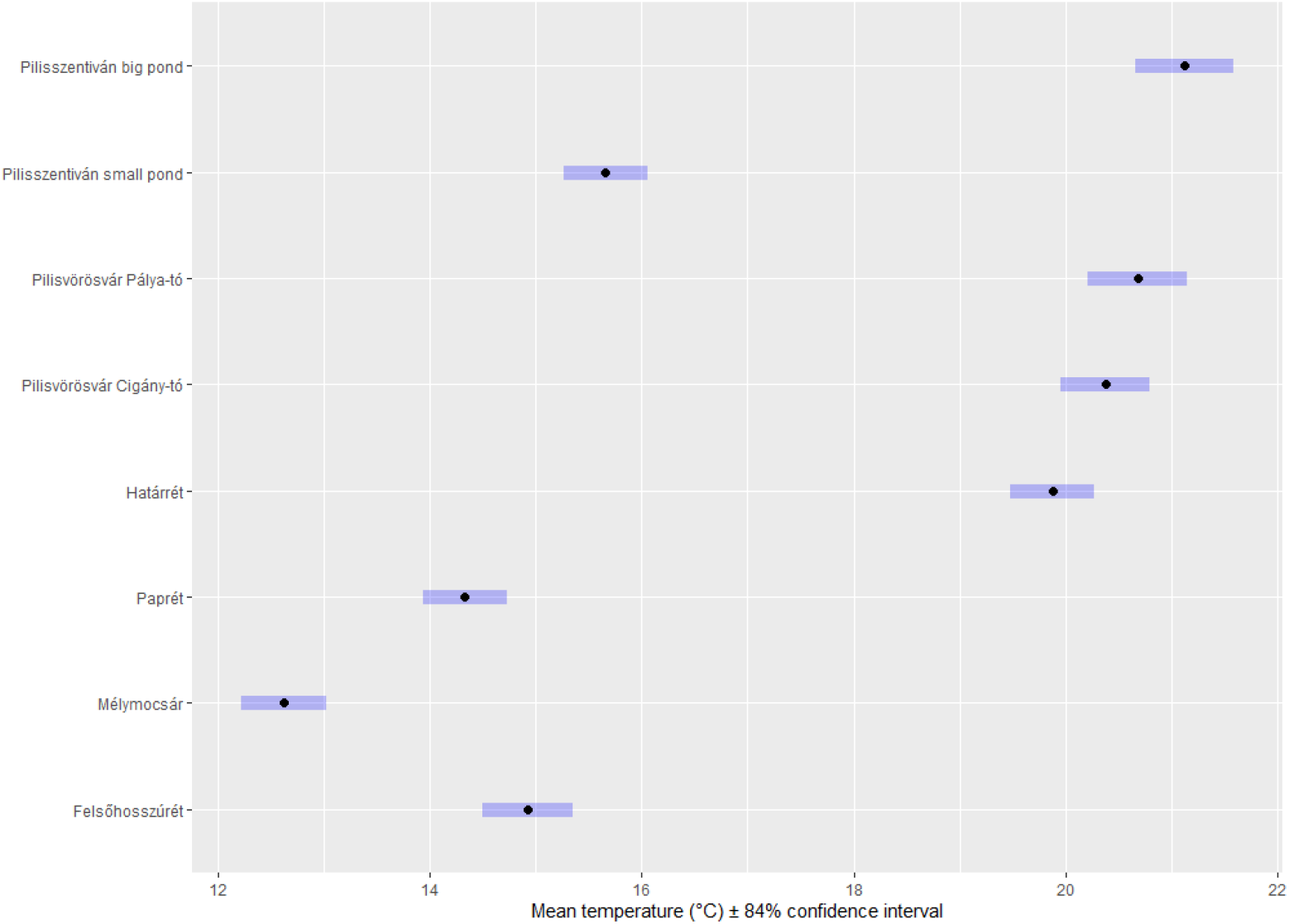
Estimated marginal means with 84 % confidence intervals for water temperature in each pond in 2022. The estimates were obtained from a generalized additive mixed model applied to the data shown in Figure S1. Non-overlapping error bars represent significant differences. Note that the smallest difference for woodland–urban pairwise comparisons was between Felsőhosszúrét and Pilisszentiván small pond (0.73 ± 0.29 °C, P = 0.016 after correction for false discovery rate).

**Figure S4.**
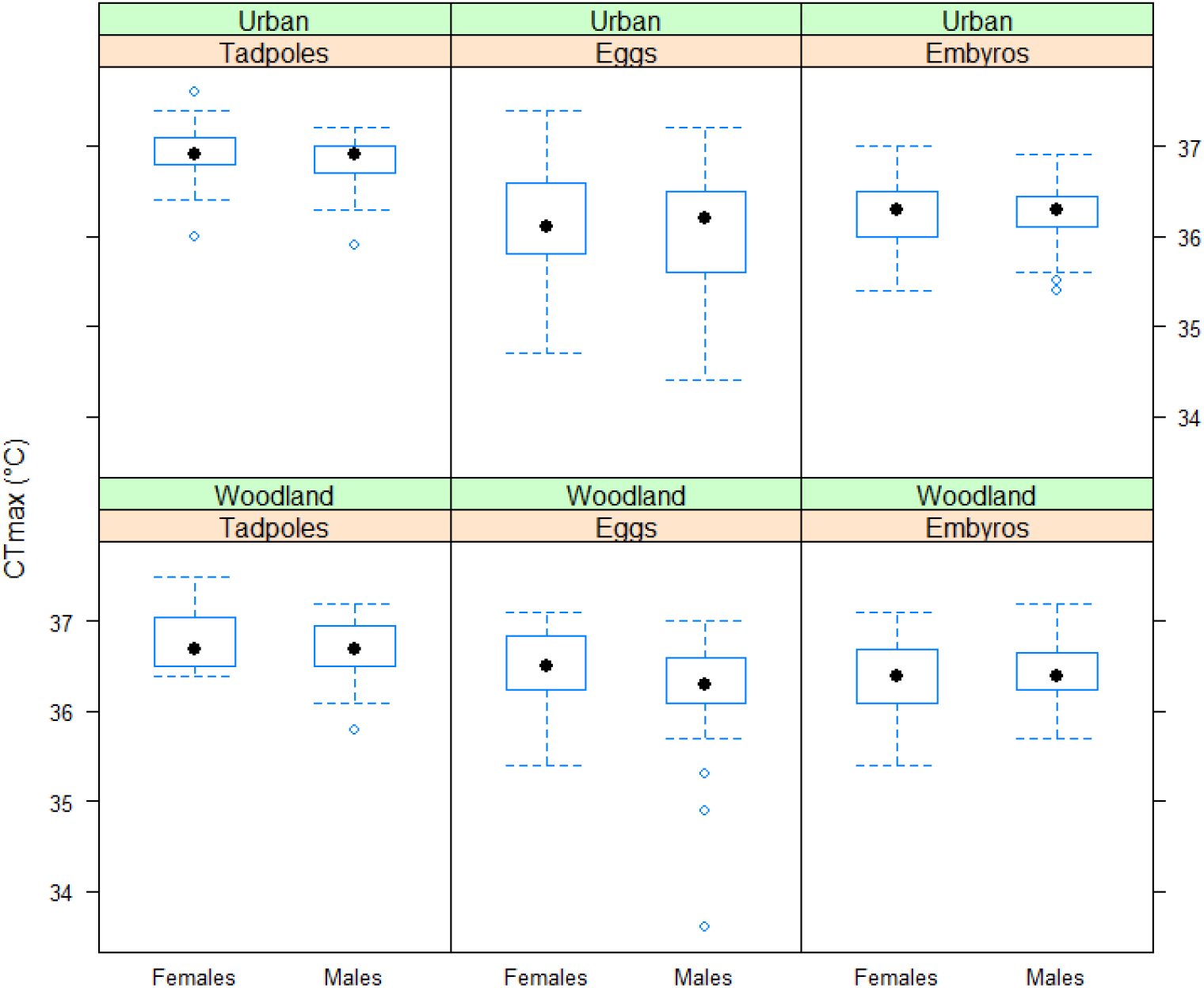
Critical thermal maximum (CT_max_) of female and male agile frog tadpoles originating from woodland or urban habitats and raised free in the field (experiencing the temperatures shown in Fig. S1) or in captivity starting from the egg or embryo stage. In each box plot, the black dot and the box represent the median and the interquartile range, respectively; the whiskers extend to the most extreme data points within 1.5 × interquartile range from the box, and the blue circles are the data points falling outside of the latter range.

## References

Åsheim, E., Andreassen, A., Morgan, R. & Jutfelt, F. 2020. Rapid-warming tolerance correlates with tolerance to slow warming but not growth at non-optimal temperatures in zebrafish. J. Exp. Biol. 223: jeb229195.

Beaman, J.E., White, C.R. & Seebacher, F. 2016. Evolution of plasticity: mechanistic link between development and reversible acclimation. Trends Ecol. Evol. 31: 237–249.

Bókony, V., Kalina, C., Ujhegyi, N., Mikó, Z., Lefler, K.K., Vili, N., et al. 2023. Does stress make males? An experiment on the role of glucocorticoids in anuran sex reversal. bioRxiv, doi: 10.1101/2023.05.27.541969.

Bókony, V., Pipoly, I., Szabó, K., Preiszner, B., Vincze, E., Papp, S., et al. 2017. Innovative females are more promiscuous in great tits (Parus major). Behav. Ecol. 28: 579–588.

Bókony, V., Ujhegyi, N., Hamow, K., Bosch, J., Thumsová, B., Vörös, J., et al. 2021a. Stressed tadpoles mount more efficient glucocorticoid negative feedback in anthropogenic habitats due to phenotypic plasticity. Sci. Total Environ. 753: 141896.

Bókony, V., Ujhegyi, N., Mikó, Z., Erös, R., Hettyey, A., Vili, N., et al. 2021b. Sex reversal and performance in fitness-related traits during early life in agile frogs. Front. Ecol. Evol. 9: 745752.

Bókony, V., Üveges, B., Ujhegyi, N., Verebélyi, V., Nemesházi, E., Csíkvári, O., et al. 2018. Endocrine disruptors in breeding ponds and reproductive health of toads in agricultural, urban and natural landscapes. Sci. Total Environ. 634: 1335–1345.

Bókony, V., Üveges, B., Verebélyi, V., Ujhegyi, N. &Móricz, Á.M. 2019. Toads phenotypically adjust their chemical defences to anthropogenic habitat change. Sci. Rep. 9: 3163.

Bonier, F., Martin, P.R., Sheldon, K.S., Jensen, J.P., Foltz, S.L. & Wingfield, J.C. 2007. Sexspecific consequences of life in the city. Behav. Ecol. 18: 121–129.

Brans, K.I., Engelen, J.M.T., Souffreau, C. & De Meester, L. 2018. Urban hot-tubs: Local urbanization has profound effects on average and extreme temperatures in ponds. Landsc. Urban Plan. 176: 22–29.

Brans, K.I., Jansen, M., Vanoverbeke, J., Tüzün, N., Stoks, R. & De Meester, L. 2017. The heat is on: Genetic adaptation to urbanization mediated by thermal tolerance and body size. Glob. Chang. Biol. 23: 5218–5227.

Campbell-Staton, S.C., Winchell, K.M., Rochette, N.C., Fredette, J., Maayan, I., Schweizer, R.M., et al. 2020. Parallel selection on thermal physiology facilitates repeated adaptation of city lizards to urban heat islands. Nat. Ecol. Evol. 4: 652–658.

Cicort-Lucaciu, A.-S., Sas, I., Roxin, M., Badar, L. & Goilean, C. 2011. The feeding study of a Rana dalmatina population from Carei Plain. South West. J. Hortic. Biol. Environ. 2: 35–46.

Diamond, S.E., Chick, L., Perez, A., Strickler, S.A. & Martin, R.A. 2017. Rapid evolution of ant thermal tolerance across an urban-rural temperature cline. Biol. J. Linn. Soc. 121: 248–257.

Diamond, S.E. & Martin, R.A. 2021. Physiological adaptation to cities as a proxy to forecast global-scale responses to climate change. J. Exp. Biol. 224: jeb229336.

Edmands, S. 2021. Sex ratios in a warming world: thermal effects on sex-biased survival, sex determination, and sex reversal. J. Hered. 112: 155–164.

Gosner, K.L. 1960. A simplified table for staging anuran embryos and larvae with notes on identification. Herpetologica 16: 183–190.

Goulet, C.T., Thompson, M.B. & Chapple, D.G. 2017a. Repeatability and correlation of physiological traits: Do ectotherms have a “thermal type”? Ecol. Evol. 7: 710–719.

Goulet, C.T., Thompson, M.B., Michelangeli, M., Wong, B.M. & Chappele, D.G. 2017b. Thermal physiology: A new dimension of the pace□of□life syndrome. J. Anim. Ecol. 86: 1269–1280.

Gunderson, A.R. & Stillman, J.H. 2015. Plasticity in thermal tolerance has limited potential to buffer ectotherms from global warming. Proc. R. Soc. B Biol. Sci. 282.

Hitchings, S.P. & Beebee, T.J.C. 1997. Genetic substructuring as a result of barriers to gene flow in urban Rana temporaria (common frog) populations: Implications for biodiversity conservation. Heredity 79: 117–127.

Hutchison, V.H. & Maness, J.D. 1979. The role of behavior in temperature acclimation and tolerance in ectotherms. Integr. Comp. Biol. 19: 367–384.

Kaya, U., Kuzmin, S., Sparreboom, M., Ugurtas, I.H., Tarkhnishvili, D., Anderson, S., et al. 2009. Rana dalmatina. IUCN Red List Threat. Species e.T58584A11790570.

Kroeker, K.J. & Sanford, E. 2022. Ecological leverage points: species interactions amplify the physiological effects of global environmental change in the ocean. Ann. Rev. Mar. Sci. 14: 75–103.

Lambert, M.R., Smylie, M.S., Roman, A.J., Freidenburg, L.K. & Skelly, D.K. 2018. Sexual and somatic development of wood frog tadpoles along a thermal gradient. J. Exp. Zool. Part A Ecol. Integr. Physiol. 329: 72–79.

Layne, J.R. & Claussen, D.L. 1982. The time courses of CTMax and CTMin acclimation in the salamander Desmognathus fuscus. J. Therm. Biol. 7: 139–141.

Leith, N.T., Fowler-Finn, K.D. & Moore, M.P. 2022. Evolutionary interactions between thermal ecology and sexual selection. Ecol. Lett. 25: 1919–1936.

Lesbarrères, D., Primmer, C.R., Lodé, T. &Merilä, J. 2006. The effects of 20 years of highway presence on the genetic structure of Rana dalmatina populations. Ecoscience 13: 531–538.

Li, D. & Bou-Zeid, E. 2013. Synergistic interactions between urban heat islands and heat waves: The impact in cities is larger than the sum of its parts. J. Appl. Meteorol. Climatol. 52: 2051–2064.

Lindauer, A.L., Maier, P.A. & Voyles, J. 2020. Daily fluctuating temperatures decrease growth and reproduction rate of a lethal amphibian fungal pathogen in culture. BMC Ecol. 20: 18.

Martin, R.A., Chick, L.D., Garvin, M.L. & Diamond, S.E. 2021. In a nutshell, a reciprocal transplant experiment reveals local adaptation and fitness trade-offs in response to urban evolution in an acorn-dwelling ant. Evolution 75: 876–887.

McLean, M.A., Angilletta, M.J. & Williams, K.S. 2005. If you can’t stand the heat, stay out of the city: Thermal reaction norms of chitinolytic fungi in an urban heat island. J. Therm. Biol. 30: 384–391.

Mitchell, N.J. & Janzen, F.J. 2010. Temperature-dependent sex determination and contemporary climate change. Sex. Dev. 4: 129–140.

Murren, C.J., Auld, J.R., Callahan, H., Ghalambor, C.K., Handelsman, C.A., Heskel, M.A., et al. 2015. Constraints on the evolution of phenotypic plasticity: Limits and costs of phenotype and plasticity. Heredity 115: 293–301.

Nemesházi, E., Gál, Z., Ujhegyi, N., Verebélyi, V., Mikó, Z., Üveges, B., et al. 2020. Novel genetic sex markers reveal high frequency of sex reversal in wild populations of the agile frog (Rana dalmatina) associated with anthropogenic land use. Mol. Ecol. 29: 3607–3621.

Nemesházi, E., Kövér, S. &Bókony, V. 2021. Evolutionary and demographic consequences of temperature-induced masculinization under climate warming: the effects of mate choice. BMC Ecol. Evol. 21: 16.

Pagliaro, M.D. & Knouft, J.H. 2020. Differential effects of the urban heat island on thermal responses of freshwater fishes from unmanaged and managed systems. Sci. Total Environ. 723: 138084.

Perkins-Kirkpatrick, S.E. & Lewis, S.C. 2020. Increasing trends in regional heatwaves. Nat. Commun. 11: 1–8.

Pike, N. 2011. Using false discovery rates for multiple comparisons in ecology and evolution. Methods Ecol. Evol. 2: 278–282.

Pottier, P., Burke, S., Drobniak, S.M., Lagisz, M. & Nakagawa, S. 2021. Sexual (in)equality? A meta-analysis of sex differences in thermal acclimation capacity across ectotherms. Funct. Ecol. 35: 2663–2678.

Preiszner, B., Papp, S., Pipoly, I., Seress, G., Vincze, E., Liker, A., et al. 2017. Problemsolving performance and reproductive success of great tits in urban and forest habitats. Anim. Cogn. 20: 53–63.

R Core Team. 2022. R: A language and environment for statistical computing. Version 4.2.2.

R Foundation for Statistical Computing, Vienna, Austria. http://www.r-project.org.

Radchuk, V., Reed, T., Teplitsky, C., van de Pol, M., Charmantier, A., Hassall, C., et al. 2019. Adaptive responses of animals to climate change are most likely insufficient. Nat. Commun. 10: 3109.

Riis, N. 1997. Field studies on the ecology of agile frog in Denmark. In: Der Springfrosch (Rana dalmatina). Ökologie und Bestandssituation, Rana Sonderheft 2 (A. Krone, K. D. Kühnel, &H. Berger, eds), pp. 189–202.

Roeder, K.A., Roeder, D. V. & Bujan, J. 2021. Ant thermal tolerance: a review of methods, hypotheses, and sources of variation. Ann. Entomol. Soc. Am. 114: 459–469.

Rohr, J.R., Civitello, D.J., Cohen, J.M., Roznik, E.A., Sinervo, B. & Dell, A.I. 2018. The complex drivers of thermal acclimation and breadth in ectotherms. Ecol. Lett. 21: 1425–1439.

Rubalcaba, J.G., Gouveia, S.F. &Olalla-Tárraga, M.A. 2019. A mechanistic model to scale up biophysical processes into geographical size gradients in ectotherms. Glob. Ecol. Biogeogr. 28: 793–803.

Ruckstuhl, K.E. & Neuhaus, P. 2005. Sexual segregation in vertebrates. Ecology of the two sexes. Cambridge University Press.

Schacht, R., Beissinger, S.R., Wedekind, C., Jennions, M.D., Geffroy, B., Liker, A., et al. 2022. Adult sex ratios: causes of variation and implications for animal and human societies. Commun. Biol. 5: 1–16.

Sykes, B.E., Hutton, P. & McGraw, K.J. 2021. Sex-specific relationships between urbanization, parasitism, and plumage coloration in house finches. Curr. Zool. 67: 237–244.

Troia, M.J. 2023. Magnitude–duration relationships of physiological sensitivity and environmental exposure improve climate change vulnerability assessments. Ecography 2023: 1–14.

Turriago, J.L., Tejedo, M., Hoyos, J.M., Camacho, A. & Bernal, M.H. 2023. The time course of acclimation of critical thermal maxima is modulated by the magnitude of temperature change and thermal daily fluctuations. J. Therm. Biol. 114: 103545.

Ujszegi, J., Bertalan, R., Ujhegyi, N., Verebélyi, V., Nemesházi, E., Mikó, Z., et al. 2022. “Heat waves” experienced during larval life have species-specific consequences on lifehistory traits and sexual development in anuran amphibians. Sci. Total Environ. 835: 155297.

Urban, M.C., Richardson, J.L. & Freidenfelds, N.A. 2014. Plasticity and genetic adaptation mediate amphibian and reptile responses to climate change. Evol. Appl. 7: 88–103.

van Steen, Y., Ntarladima, A.M., Grobbee, R., Karssenberg, D. & Vaartjes, I. 2019. Sex differences in mortality after heat waves: are elderly women at higher risk? Int. Arch. Occup. Environ. Health 92: 37–48.

Vences, M., Hauswaldt, J.S., Steinfartz, S., Rupp, O., Goesmann, A., Künzel, S., et al. 2013. Radically different phylogeographies and patterns of genetic variation in two European brown frogs, genus Rana. Mol. Phylogenet. Evol. 68: 657–670.

Zuur, A.F., Ieno, E.N. J. Walker, N., Saveliev, A.A. & Smith, G.M. 2009. Mixed effects models and extensions in ecology with R.Springer, New York.

